# Transcutaneous auricular Vagus nerve stimulation for working memory enhancement: A comparative study of electrical and ultrasound stimulation

**DOI:** 10.64898/2026.03.06.710106

**Authors:** Alicia Falcon-Caro, Nicholas Myers, Marcus Kaiser, Hyuk Choi, Jae-Jun Song, JeYoung Jung

## Abstract

**Objectives:** Transcutaneous auricular vagus nerve stimulation (taVNS) is a non-invasive neuromodulation technique that has shown potential to enhance cognitive function, including working memory. This study investigated the acute effects of both electrical (E-taVNS) and ultrasound (U-taVNS) modalities on working memory using a 3-back task in healthy young adults. We hypothesized that active taVNS would enhance working memory performance relative to sham, and that both stimulation modalities would engage similar neuromodulatory mechanisms.

**Materials and Methods:** Fifty-nine participants underwent a single-blind, sham-controlled, within-subject design study, with working memory performance assessed using a 3-back task before and after stimulation. Primary performance measures included correct rejection rate, error false alarm, and sensitivity (*d*′). Statistical analyses compared pre- and post-stimulation performance across modalities.

**Results:** E-taVNS significantly enhanced working memory performance through an increase in correct rejection rate and sensitivity (measured by *d’*), alongside a reduction in error of false alarm. U-taVNS showed a similar directional trend across performance measures, although these effects did not reach statistical significance. Baseline anxiety levels significantly predicted individual responsiveness to taVNS. In terms of tolerability, a higher proportion of participants receiving E-taVNS reported skin irritation compared to those receiving U-taVNS.

**Conclusions:** E-taVNS can acutely enhance working memory performance, while U-taVNS may offer a comparable, better tolerated alternative. Our findings highlight the potential of taVNS to support memory function, while showing the importance of further research to clarify modality-specific effects and optimize stimulation parameters.

## Introduction

Working memory (WM) is a core cognitive system responsible for the temporary storage and manipulation of information required for higher-order cognition, including reasoning, learning, and decision-making [1, 2]. WM capacity varies across individuals and can predict differences in attentional control, fluid intelligence, and complex cognitive performance [3]. Furthermore, impairments in WM are well-established markers of several neurological and psychiatric disorders, including attention-deficit/hyperactivity disorder (ADHD) [4], schizophrenia [5], traumatic brain injury [6], and neurodegenerative conditions, such as Alzheimer’s disease [7, 8]. WM is a dynamic process that can be modulated by neuromodulatory pathways through the regulation of noradrenergic input from the locus coeruleus (LC) and cholinergic projections from the basal forebrain [9–12]. These neuromodulatory systems are thought to enhance signal-to-noise ratio of task-relevant representations, especially under high cognitive demand [13]. Therefore, WM represents a promising target for neuromodulation approaches aimed at enhancing cognitive function in both healthy and clinical populations at risk of cognitive decline [14].

Behavioral and non-invasive brain stimulation approaches, including Transcranial Magnetic Stimulation (TMS) and transcranial Electrical Stimulation (tES) have targeted memory networks to mitigate cognitive decline [15, 16]. While some studies reported positive outcomes following cortical stimulation, others yielded inconsistent findings. In contrast, transcutaneous auricular Vagus Nerve Stimulation (taVNS) modulates cognitive performance processes indirectly through vagal pathways rather than through direct modulation of cortical excitability and has shown promising effects on working memory enhancement [17, 18].

Among emerging neuromodulation techniques, Vagus Nerve stimulation (VNS) has received increasing attention due to its capacity to influence large-scale brain networks involved in cognitive control. The Vagus Nerve consists predominantly of afferent fibers that convey sensory information to the nucleus tractus solitarius (NTS) in the brainstem, which in turn projects to neuromodulatory centers including the LC, hippocampus, and prefrontal cortex [19–21]. Through these pathways, VNS can modulate arousal, attention, and executive function [22].

While traditional invasive VNS requires surgical implantation, which is associated with procedural risks and limited scalability [23], taVNS offers a non-invasive alternative by stimulating the auricular branch of the Vagus Nerve (ABVN) at the external ear, usually at the cymba conchae or tragus [23].

Furthermore, neuroimaging studies have shown that taVNS can engage central vagal pathways, including the NTS and LC, with activation patterns resembling those observed during invasive VNS [20, 24]. These findings suggest that taVNS can access neuromodulatory systems relevant to cognitive control without surgical intervention. As discussed in [21] and [25], taVNS is proposed to affect WM through the modulation of LC-norepinephrine (LC-NE) system. Noradrenergic input to the prefrontal cortex is also known to support WM by enhancing task-relevant representations and suppressing distraction [26]. Consistent with this, taVNS has been associated with physiological markers of LC–NE engagement, including pupil dilation [27, 28], P300 event-related potentials [29], and salivary alpha-amylase [30, 31]. Together, these findings suggest that taVNS may improve WM by optimizing neuromodulatory tone rather than directly altering local cortical excitability. Despite this mechanistic rationale, empirical evidence for taVNS effects on WM remains mixed. Some studies report improvements in reaction time or efficiency, particularly when stimulation is applied offline [18, 32], while others observe limited or task-specific effects on accuracy and sensitivity [33–35]. Such inconsistencies likely reflect methodological heterogeneity across studies, including differences in stimulation parameters, timing, and WM tasks. Moreover, despite the potential as a relevant intervention for neurodegenerative diseases, relatively few studies have examined taVNS effects under high cognitive load, such as demanding n-back tasks. Traditionally, taVNS has been predominantly applied using electrical stimulation. However, recent advances in neuromodulation have introduced low-intensity transcranial focused ultrasound (FUS) as a promising non-invasive method. This neuromodulation technique has also emerged as a novel non-invasive approach for VNS [36, 37], including ABVN [38–40].

Unlike traditional electrical taVNS (E-taVNS), ultrasound neurostimulation operates through mechanical energy, inducing pressure changes in neural membranes that can elicit action potential propagation without cutaneous discomfort [41]. Preliminary evidence suggests that ultrasound taVNS (U-taVNS) may modulate autonomic and affective processes [39], potentially engaging central vagal pathways relevant to cognition similarly to E-taVNS while offering improved tolerability.

Even so, no studies to date have directly compared electrical and ultrasound taVNS within the same experimental framework. As a result, it remains unclear whether these two non-invasive stimulation modalities exert comparable effects on working memory or differ in efficacy and sensitivity to individual differences.

Therefore, in the present study, we directly compared the effects of electrical and ultrasound taVNS on working memory performance in healthy adults using a high-load 3-back task. Participants received either E-taVNS or U-taVNS under active and sham conditions and completed the 3-back task before and after stimulation. Individual differences in anxiety, depression, and interoceptive perception were evaluated to examine variability in stimulation responsiveness. We hypothesized that active taVNS would enhance working memory performance relative to sham, particularly under high cognitive load, and that both stimulation modalities would engage similar neuromodulatory mechanisms. Furthermore, we compared the efficacy of E-taVNS and U-taVNS and explored whether baseline traits were associated with individual variability in taVNS effects.

## Materials and Methods

### Participants

An a-priori sample size calculation was performed using G*Power 3.1.9.7 [42] to estimate the approximate number of participants required for a within-subject design (power= 0.8, alpha= 0.05). Based on a moderate effect size based on previous study [18], the estimated sample size was 19 participants. To ensure sufficient power and allow for potential attrition, we aimed to recruit 30 participants per stimulation modality.

59 healthy adults (46 females, 13 males; *M* age = 23.6, SD=2.88) were recruited and randomly allocated to either the electrical taVNS group (E-taVNS; *n* = 30) or the ultrasound taVNS group (U-taVNS; *n* = 29). Eligibility was screened using the VNS Safety Questionnaire. Exclusion criteria included: age *<* 18 years; active implantable medical devices (e.g., pacemaker, cochlear implant); history of carotid atherosclerosis or cervical vagotomy; cardiovascular conditions (e.g., hypertension, hypotension, bradycardia, tachycardia); metallic implants in the head or neck; current or past neurological or psychiatric disorders; predisposition to fainting; family history of epilepsy; current use of psychoactive medication (except hormonal contraceptives); and pregnancy. These criteria ensured participant safety and data integrity. Due to missing data from Sham session, one female participant from U-taVNS was excluded from the analysis.

All participants provided written informed consent. The study was approved by the University of Nottingham Research Ethics Committee (F1619R) and conducted in accordance with the Declaration of Helsinki.

### Experimental Design and Procedures

Participants completed the Beck Anxiety Inventory (BAI; [43]), the Beck Depression Inventory (BDI; [44]), and the Multidimensional Assessment of Interoceptive Awareness (MAIA-2; [45]) before taking part in the experiment.

Each participant completed two sessions: one active stimulation session and one sham session, separated by at least five days. Session order was counterbalanced across participants. At the start of each session, participants completed a 3-back working memory task. Following these baseline assessments, participants received 30 minutes of taVNS. Immediately after stimulation, the task was repeated (Fig. 1a). At the end of each session, participants completed a side-effects questionnaire assessing mild aversive sensations (e.g., headache, nausea, neck tension, tingling).

**Figure 1.**
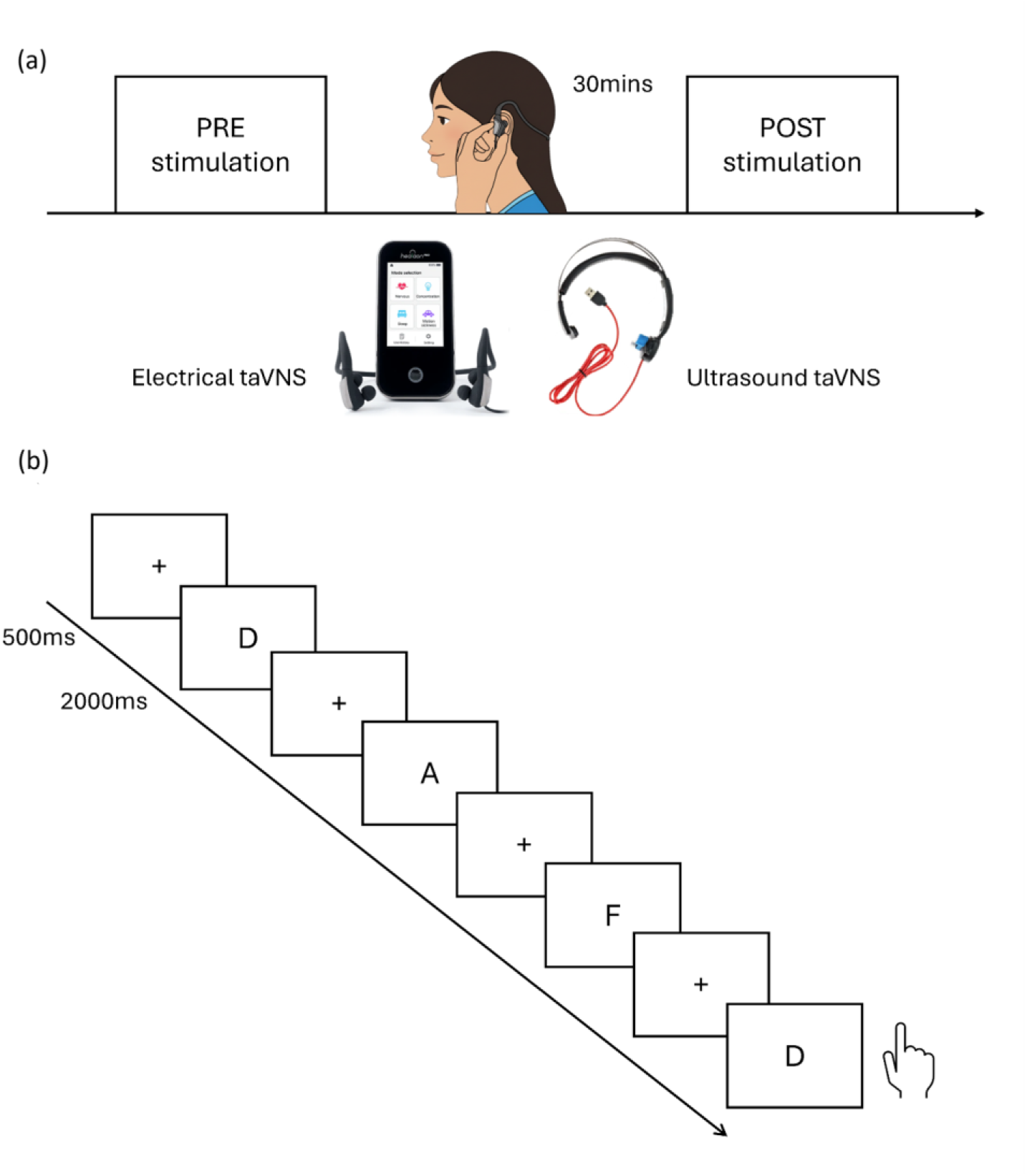
Experimental procedure and design. (a) Schematic overview of the experimental procedure. Each participant completed two sessions (active and sham), separated by at least five days. At the start of each session, participants performed a working memory task (pre-stimulation). Participants then received 30 minutes of transcutaneous auricular Vagus nerve stimulation (taVNS) using either electrical (E-taVNS) or ultrasound (U-taVNS) stimulation, depending on group assignment. Immediately following stimulation, the task was repeated (post-stimulation). (b) Trial structure of the 3-back working memory task. On each trial, an uppercase letter was presented at the center of the screen for 2 seconds, followed by a 0.5 seconds fixation cross. Participants responded by pressing the target key when the current letter matched the letter presented three trials earlier; all other letters required withholding a response.

Working memory was assessed using a 3-back task programmed in PsychoPy (version 2024.2.4) (Fig. 1b). On each trial, an uppercase letter was presented for 2 seconds followed by a 0.5 seconds fixation cross. Participants pressed the spacebar when the current letter matched the letter from three trials earlier; all other trials required a non-response. The task comprised 50 trials, with 33% designated as targets. A brief practice block was administered prior to the task. Procedures were consistent across sessions and stimulation modalities.

### Transcutaneous auricular Vagus Nerve Stimulation (taVNS)

Two stimulation modalities were used: electrical taVNS (E-taVNS) and ultrasound taVNS (U-taVNS) (Fig. 1a bottom).

Electrical stimulation was administered bilaterally to the cymba conchae using conductive rubber electrodes (Healaon Pro, Neurive Inc., Gimhae, Republic of Korea). Stimulation parameters were 30 Hz frequency and level 5 intensity (2.5 mA). Each session lasted 30 minutes. A continuous wind-like auditory background masked device noise in both active and sham conditions. Sham stimulation consisted of 30 seconds of stimulation at the beginning of the session, after which no further current was delivered; device placement and auditory masking were identical to the active condition.

Ultrasound stimulation was delivered to the right cymba conchae using the ZenBud device (NeurGear Inc., Rochester, NY, USA). Parameters included a 5.3 MHz center frequency, 41 Hz pulse repetition frequency, 50% duty cycle, peak mechanical output of 0.081 W/cm^2^, and average intensity of 1.03 MPa. Sham stimulation was administered using an identical sham device that delivered no active ultrasound.

### Statistical Analysis

All statistical analyses were performed in Python (version 3.13). To quantify stimulation-related changes in working memory performance (overall accuracy, hit accuracy, correct rejection rate, average hit reaction time, and *d’*) as Post/Pre ratio metrics were computed per participant. Ratio metrics were used to normalize post-stimulation performance to individual baseline differences.

To comprehensively evaluate the main effects of modality (E-taVNS vs U-taVNS), session (Pre vs Post), and stimulation (Active vs Sham), and their interactions, a Linear Mixed Effects Model (LMEM) was implemented for each performance metric. This LMEM (a 2×2×2 mixed design) was the primary test for differential treatment effects. Post-hoc pairwise comparisons assessed changes between pre-, and post-stimulation phases. To complement this analysis and account for practice effects and inter-subject variability in baseline performance, a baseline-adjusted LMEM was also conducted, using post-stimulation values as the dependent variable and pre-stimulation performance as covariate. This approach provides an estimate of stimulation effects while mitigating regression to the mean and validating the primary LMEM results.

To examine modality effects (E-taVNS vs U-taVNS) and stimulation (Active vs Sham), a two-way mixed-design ANOVA was performed on the Post/Pre ratio metrics, with stimulation condition as a within-subject factor and modality as a between-subject factor.

To confirm no significant difference in baseline between modalities, an independent samples Student’s t-test was conducted on all reported metrics.

To evaluate direct taVNS effects, paired t-tests were performed on the Post/Pre ratio outcomes between active and sham conditions for both modalities (E-taVNS and U-taVNS) separately, directly assessing taVNS effects within each modality. Effect sizes were estimated using Cohen’s *d*.

Visual inspection of the distributions (Fig. 2 and 3) suggested the presence of at least one potential outlier. Therefore, outliers were identified using the interquartile (IQR) criterion. Analyses were repeated after excluding these observations. However, the statistical conclusions remained unchanged. Therefore, the results reported in the main analysis include the outliers.

**Figure 2.**
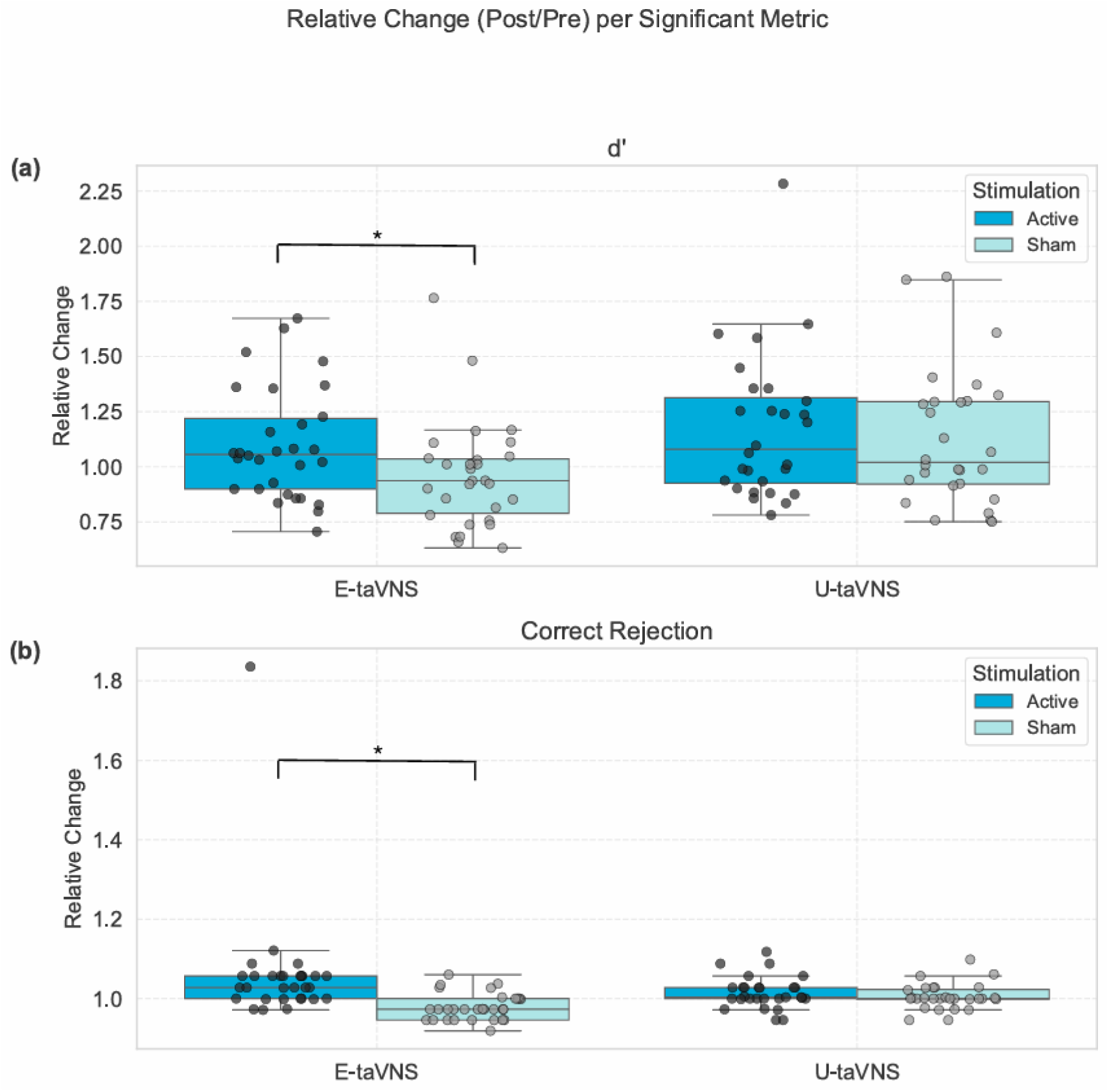
Effects of electrical and ultrasound taVNS on working memory. The two main metrics which showed significant effect of condition (active vs sham) are included. (a) presents *d*′ ratio (post/pre) for E-taVNS and U-taVNS separately under active and sham stimulation, while (b) presents results for correct rejection ratio. Each dot represents individual participants per modality.

**Figure 3.**
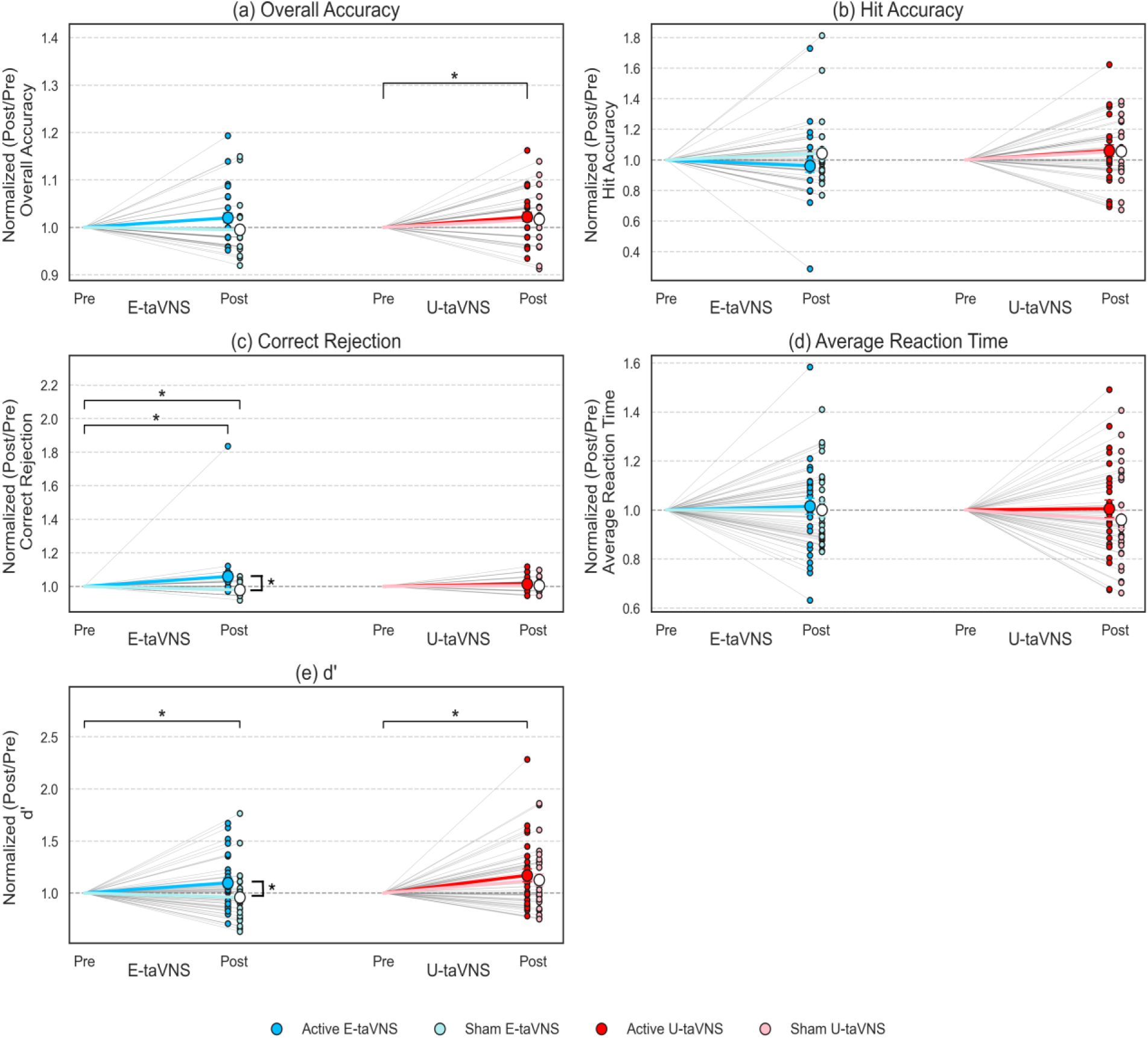
Individual and group-level changes in working memory performance from pre- to post-stimulation. Pre- to post-stimulation Post/Pre ratio changes in working memory metrics are shown for E-taVNS and U-taVNS under active and sham conditions. Each panel (a-e) corresponds to a different metric (Overall Accuracy, Hit Accuracy, Correct Rejection rate, Average Hit Reaction Time, and *d*′). For each modality, pre-stimulation values are normalized to 1 and shared across stimulation conditions. Thin grey lines represent individual participant trajectories from pre- to post-stimulation, with colored points indicating post-stimulation values for Active and Sham conditions. Thick colored lines depict group means, with error bars indicating SD. Colors denote stimulation condition within each modality (Active vs Sham), allowing visual comparison of stimulation-specific effects within and across modalities. Significant differences (p<0.05) are represented by brackets and ‘*’ for Post vs Pre, and Active vs Sham ratio.

To examine aversive effects, a series of chi-square tests were conducted to compare frequencies of reported side effects across stimulation conditions (Active vs Sham) and modality (E-taVNS vs U-taVNS). All results are considered significant if *p <* 0.05.

Finally, to identify baseline traits associated with stimulation responsiveness, Pearson Correlation analysis was performed between baseline questionnaire scores and the efficacy ratio (Active Post/Pre divided by Sham Post/Pre). This ratio served as the dependent variable to isolate the neuromodulation effect from baseline differences. Linear regression analysis was used to further characterize any significant predictive relationships.

## Results

### Demographics Results

Independent-sample t-tests and chi-square t-tests were conducted on demographic and psychological variables to ensure that the two groups, E-taVNS and U-taVNS, were comparable prior to experimental manipulation. No significant differences between modalities in sex distribution (Female/Male: E-taVNS: 23/7, U-taVNS: 23/6; *X*^2^ = 0.00; *p* = 1.00), or age (E-taVNS: 23.58 ± 2.88, U-taVNS: 23.79 ± 3.31; *t* = −0.28; *p* = 0.781) were found. Due to missing sham results, one female participant from U-taVNS was removed from the analysis. Therefore, a total of 58 participants were analyzed, 30 for E-taVNS and 28 for U-taVNS.

Furthermore, no group differences were found in anxiety (BAI: E-taVNS: 8.87 ± 6.66, U-taVNS: 9.66±6.11; *t* = −0.47; *p* = 0.637), depression (BDI: E-taVNS: 7.00±5.84, U-taVNS: 6.86±7.68; *t* = 0.08; *p* = 0.939), or MAIA (total: E-taVNS: 2.89±0.63, U-taVNS: 3.00 ± 0.52; *t* = −0.74; *p* = 0.463).

**Table 1.** Baseline Demographic and Psychological Characteristics of Participants between Modalities. Values represent mean ± standard deviation (SD). Significance difference is considered for *p <* 0.05.

|  | E-taVNS (n = 30) | U-taVNS (n = 29) | Test Statistic | p-value |
| --- | --- | --- | --- | --- |
| Sex (Female/Male) | 23 / 7 | 23 / 6 | $\chi^2 = 0.00$ | 1.000 |
| Age (years) | $23.58 \pm 2.88$ | $23.79 \pm 3.31$ | $t = -0.28$ | 0.781 |
| Anxiety (BAI) | $8.87 \pm 6.66$ | $9.66 \pm 6.11$ | $t = -0.47$ | 0.637 |
| Depression (BDI) | $7.00 \pm 5.84$ | $6.86 \pm 7.68$ | $t = 0.08$ | 0.939 |
| MAIA (Total) | $2.89 \pm 0.63$ | $3.00 \pm 0.52$ | $t = -0.74$ | 0.463 |

### Effect of taVNS on Working Memory

As there were practice effects and inter-individual variability in baseline (Table S1), we decided to use the Post/Pre ratio to be reported in the results.

ANOVA revealed that on E-taVNS, a significant effect of condition (Active vs Sham) was found on *d*’ Post/Pre ratio (*t*(30) = 2.09, *p* = 0.045, Cohen’s *d* = 0.38), and correct rejection rate ratio (*t*(30) = 2.88, *p* = 0.007, Cohen’s *d* = 0.53), as shown in Fig. 2. This effect indicates an increase in correct rejection rate, and an increase in *d’* with respect to sham condition, which can be interpreted as an increase in working memory performance during active vs sham stimulation. However, on U-taVNS, there was no significant effect of condition across any of the ratio metrics (*d’* ratio: *t* (28) = 0.50, *p* = 0.619, Cohen’s *d* = 0.09) (Fig. 2).

LMEM revealed no significant effect of Group in the interaction between Group, Condition and Phase on *d’* (B = -0.316, SE = 0.265, *z* = -1.193, *p* = 0.233), Overall accuracy (B = -0.018, SE = 0.019, *z* = -0.942, *p* = 0.346), Hit accuracy (B = 0.065, SE = 0.053, *z* = 1.225, *p* = 0.221), or average Hit reaction time (B = 0.014, SE = 0.056, *z* = 0.245, *p* = 0.807). However, there was significant effect for correct rejection (B = - 0.056, SE = 0.018, *z* = -3.077, *p* = 0.002). Baseline-adjusted LMEM confirmed the effects observed in LMEM, with comparable effect size and significance levels, demonstrating that the observed improvements are independent of baseline variability. Results reflect a similar performance on both stimulation modalities, with larger effects on E-taVNS, as can be seen in Fig. 3. Post-hoc comparisons revealed significant increases of *d’* (*t* (28) = 2.72, *p* = 0.012, Cohen’s *d* = 0.51) and Overall accuracy (*t* (28) = 2.23, *p* = 0.034, Cohen’s *d* = 0.42) from pre- to post-stimulation in U-taVNS active condition. In E-taVNS, Correct rejection in active condition showed statistical significance between pre- to post-stimulation (*t* (30) = 2.98, *p* = 0.006, Cohen’s *d* = 0.54) and sham E-taVNS on Correct rejection (*t* (30) = −3.52, *p* = 0.001, Cohen’s *d* = −0.64) and *d’* (*t*(30) = −2.11, *p* = 0.044, Cohen’s *d* = −0.38).

It is noted that results on sham U-taVNS show a similar performance to active conditions that are not present in sham E-taVNS (Fig. 3). The individual-level raw data is reported in Fig. S1 for both conditions and modalities. The statistical improvement on *d*’ and overall accuracy from pre- to post-stimulation during active U-taVNS stimulation, although not significant compared to sham results, shows that U-taVNS can improve working memory performance.

### The relationship between taVNS Effects and Questionnaires

Linear regression analyses were conducted to examine whether baseline anxiety, measured by BAI, predicted the working memory outcome, measured as efficacy ratio (Active Post/Pre divided by Sham Post/Pre), from taVNS, when considering both modalities together. When using the BAI score as a predictor, the regression model was statistically significant (B = -0.019, SE = 0.009, t = -2.17, *p* = 0.034), accounting for 7.8% of the variance in the outcome (Fig. 4). We focused on the correlation with overall taVNS instead of specific to each modality to find overlapping relationships between the modalities and the baseline questionnaires.

**Figure 4.**
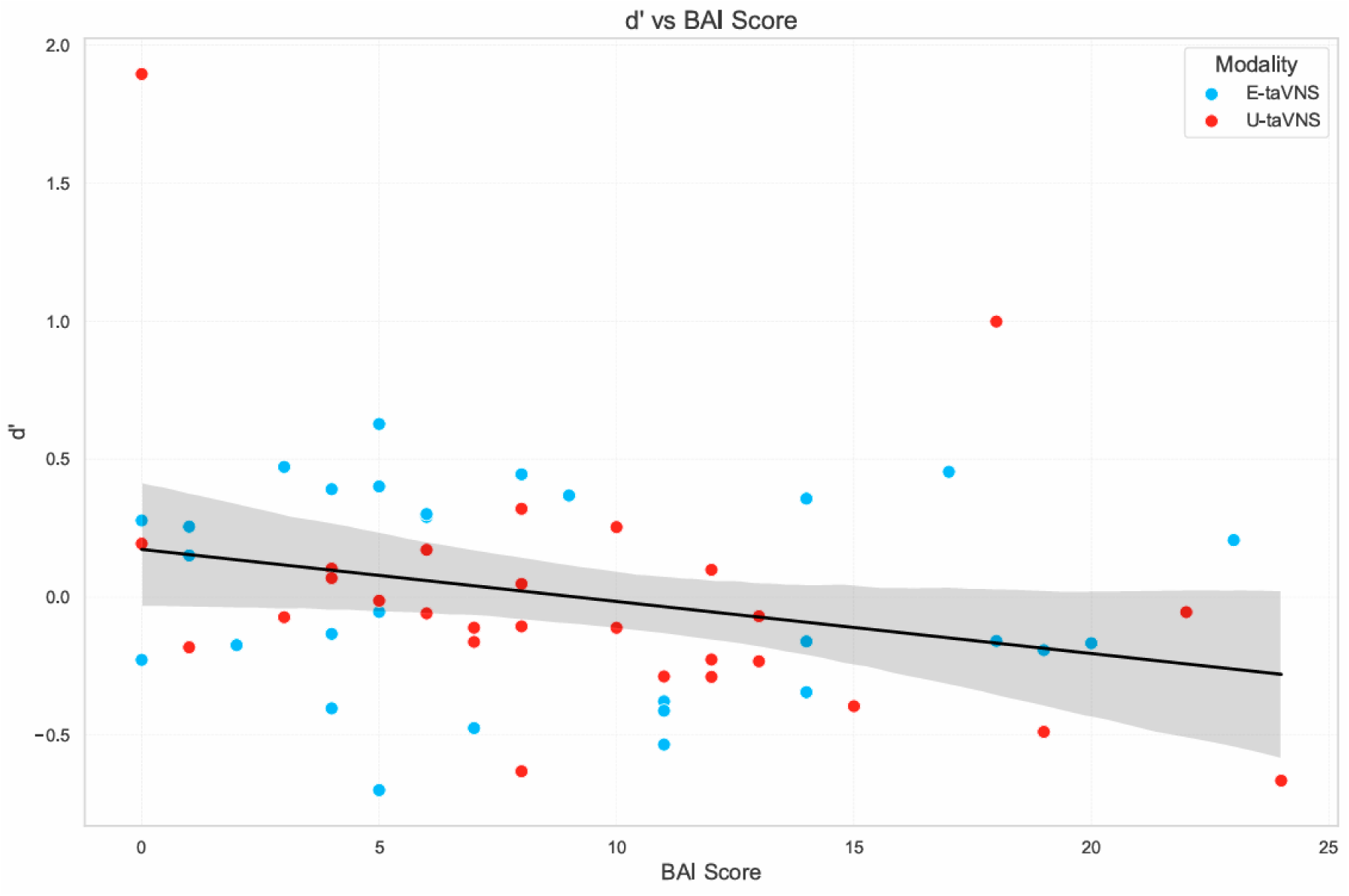
Relationship between BAI scores and outcome of taVNS in working memory performance. Scatterplot illustrates the association between BAI scores and working memory performance after taVNS, as measured by ratio changes (active/sham) in *d’*. Each point represents individual participants. Blue dots indicate participants that received E-taVNS, while red dots represent participant that received U-taVNS. A higher BAI score was associated with a worse performance and a smaller effect of the stimulation.

There were no significant correlations between the total MAIA score, BDI, and taVNS effects. Although some MAIA subscales yielded statistical significance as predictors of some working memory metrics, only BAI obtained significant results as a predictor for *d*′, which is the main metric used to assess working memory performance (Fig. S2).

When considering both modalities independently, other questionnaires also yielded statistical significance as predictors (Fig. S3). However, no questionnaire in common could predict the outcome of *d*’ for both modalities. Not worrying was highlighted as a significant predictor for Hit accuracy (Fig. S4) on E-taVNS (B = 0.102, SE = 0.043, t = 2.35, *p* = 0.026) and for Average Hit Reaction Time (Fig. S5) for U-taVNS (B = - 0.176, SE = 0.056, t = -3.15, *p* = 0.004). As shown in Fig. S3, while only not-worrying and noticing scores yield statistical significance as predictors for E-taVNS performance, for U-taVNS, multiple other baseline questionnaire metrics yield statistical significance as outcome predictors, such as BDI or Body listening. We can infer from these results that the U-taVNS outcome is more likely influenced by participants’ baseline state differences than E-taVNS.

Given that BAI and not worrying are highlighted as the baseline scores that can predict the stimulation outcome, this aligns with previous research that shows that working memory performance is influenced by attention [46] and consequently influenced by anxiety traits and stress [47].

### Comparison of Aversive Effects per Modality

A chi-square test of independence was performed to examine the association between stimulation modality and the occurrence of aversive effects (Fig. 5). A higher proportion of participants in the E-taVNS group reported skin irritation (20.0%) compared to those in the U-taVNS group (3.4%) (*X*^2^ = 3.86*, p* = 0.049), indicating U-taVNS was associated with a lower incidence of skin irritation than E-taVNS. Although other aversive effects were reported, none yielded statistical significance on the incidence between the two modalities. A portion of participants also reported ear pain (E-taVNS: 16.7%; U-taVNS: 10.3%), but no significant differences in incidence were found between the two modalities (*X*^2^ = 0.50*, p* = 0.478).

**Figure 5.**
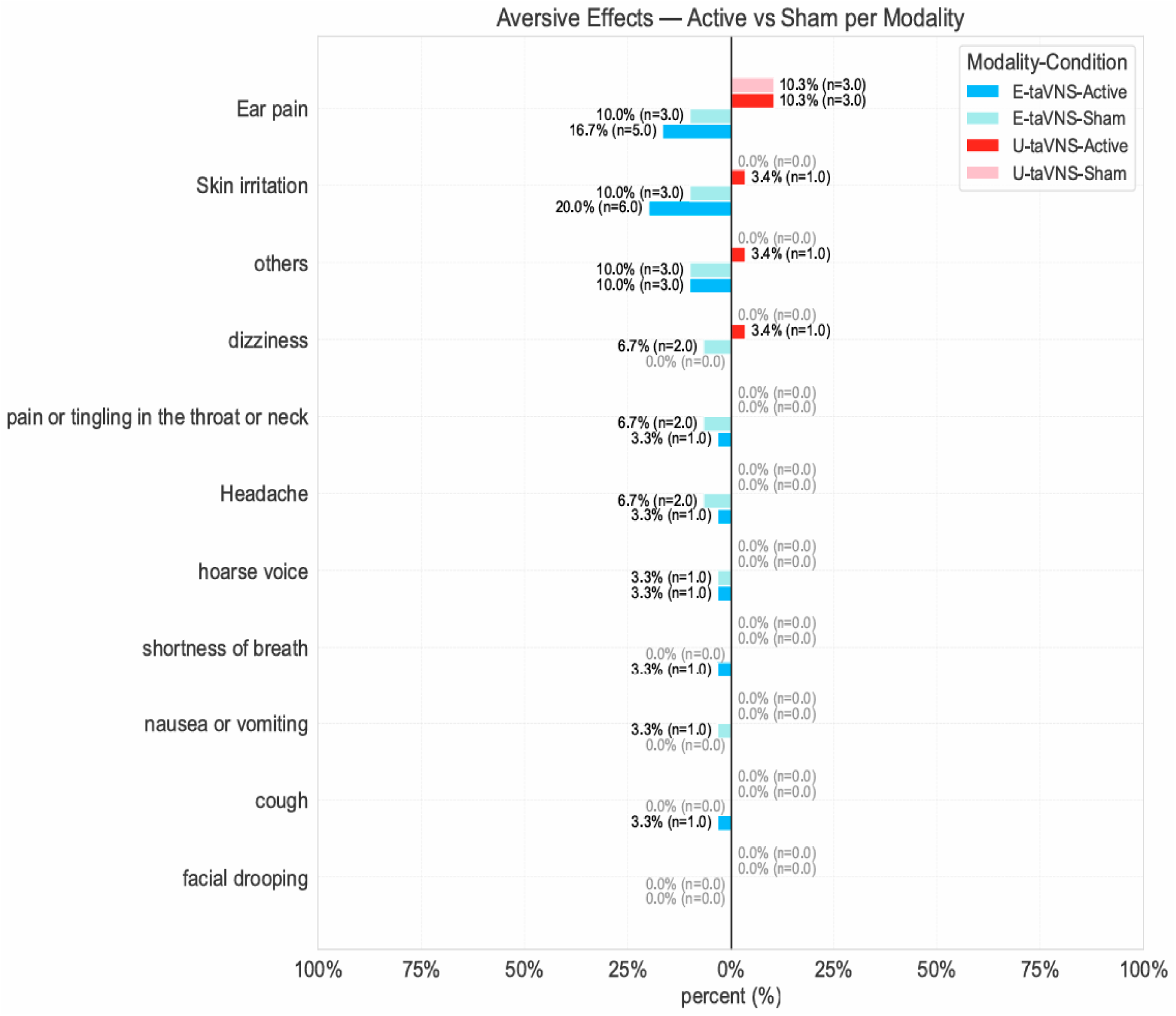
Comparison of Aversive Effect Profiles across E-taVNS and U-taVNS Modalities. Incidence of aversive effects reported by participants following E-taVNS and U-taVNS stimulation under two conditions (active and sham). The percentage and absolute count of participants reporting each aversive effect is plotted on the x-axis (e.g., *X.X*% (*n* = *Y*)). Results from E-taVNS modality (active and sham) are plotted to the left of the midline, while results for U-taVNS are plotted to the right side. Aversive effects with 0% incidence are labelled gray. This analysis directly compared the aversive effects associated with each stimulation modality and condition (active vs sham), providing key insights into the safety and tolerability of the respective stimulation modalities.

## Discussion

The present study investigated the effects of both electrical and ultrasound transcutaneous Vagus Nerve stimulation (taVNS) on working memory in healthy young adults, using a high cognitive load 3-back task.

Our findings revealed that electrical stimulation (E-taVNS) successfully improved working memory performance. While ultrasound stimulation (U-taVNS) also showed improvements in working memory performance similar to E-taVNS, the results were not significant compared to its own sham. Still, consistent with our primary hypothesis, active taVNS significantly improved working memory compared with both sham and baseline (pre-stimulation) conditions. These findings align with previous research showing that taVNS modulates higher order cognitive processing [18, 32, 48, 49]. The main effect of phase, along with post-hoc evidence of working memory improvement from pre- to post-stimulation, indicates that taVNS effects persist beyond the stimulation period.

The improvement in working memory likely reflects the modulation of multiple interconnected systems and supports the idea that working memory does not rely on a single brain network but can be influenced by a combination of other nodes and networks, as described in [50, 51]. The Vagus nerve’s afferent projections to NTS and subsequent connections to LC implicate noradrenergic mechanisms as primary mediators [21, 25]. Additionally, taVNS-induced changes in parasympathetic tone may contribute to emotional regulation [22]. Anxiety and stress levels have been proven to have an effect on memory consolidation of recent experiences and memory retrieval, which can influence individual’s performance on high-load working memory tasks, such as 3-back [47]. Therefore, the possible calming effect of taVNS could also positively influence spatial working memory performance. Although reduced anxiety is also generally beneficial in any attention-demanding tasks, this is especially relevant for high-load cognitive tasks, such as working memory, which critically depend on sustained executive and attentional control [46, 52]. This is also supported by the results presented in this paper, where BAI scores, as a measure of baseline anxiety levels, becomes a significant predictor for taVNS.

Given that taVNS effects were seen through a significant improvement in correct rejections, but not on hit accuracy, this suggests that non-targets were more effectively removed from the focus of attention and from the foreground activity subspace, as suggested in [53]. This mechanistic model, together with the results presented in this study, may suggest that the reduction in signal-to-noise (SNR) through LC originating from taVNS may indicate a reduction in noise rather than an increase in signal.

The similar trend in performance of both stimulation modalities supports the idea that activation of ABVN afferents, rather than the specific method of activation, underlies the observed effects, highlighting the importance of anatomical targeting over stimulation modality. These findings are consistent with the well-established mechanisms of taVNS action.

Although more research is needed, in addition to electrical stimulation, ultrasound-based taVNS may engage similar neural circuits through distinct biophysical mechanisms. FUS induces mechanical membrane deformation that influences neuronal excitability without direct electrical current. Given its capacity to activate the same vagal afferent pathways [38, 39], U-taVNS may produce comparable affective modulation with more compliance and tolerability.

Although significant differences in efficacy between electrical and ultrasound stimulation were found, results from ultrasound stimulation showed a similar trend to electrical stimulation. Both E-taVNS and U-taVNS produced improvements in working memory, suggesting that these distinct physical modalities may converge on shared neurophysiological pathways. This equivalence is particularly noteworthy given the fundamental differences in mechanisms, with electrical stimulation involving direct current-induced depolarization, whereas ultrasound stimulation relies on mechanical pressure waves that deform neuronal membranes and influence excitability [54]. This distinction in mechanisms can be associated with the stronger effect of electrical stimulation over ultrasound stimulation. Furthermore, an important consideration is that ultrasound in this study was applied only unilaterally. Thus, while both modalities showed similar outcomes, this comparison should be interpreted with caution, as ultrasound stimulation might have yielded stronger effects, and significant differences, had it been delivered bilaterally. This point is particularly relevant given that the electrical taVNS device used in this study stimulated both sides, whereas most conventional taVNS approaches target only the left ear due to ongoing debate about potential right-sided cardiac influences [24, 55]. Given the lack of significant difference in U-taVNS, which might be due to the difference in sham masking and delivering methods between the modalities, further research on this modality to validate this claim is still needed.

Despite the similar efficacy, U-taVNS demonstrated greater tolerability. Only 3.4% of participants in the ultrasound group reported skin irritation compared with 20.0% in the electrical group, suggesting that ultrasound stimulation may be a more tolerable alternative for long-term therapeutic applications. This improved tolerability could enhance treatment adherence, particularly important for conditions requiring extended or continuous stimulation protocols such as dementia or schizophrenia disorders. While the MAIA scores did not significantly predict the changes in working memory performance under combined taVNS, the BAI scores emerged as a significant predictor, consistent with previous studies that suggested working memory individual variability is affected by anxiety and stress conditions [47]. This finding highlights the potential relevance of anxiety in mediating taVNS effects on cognitive processing. On the other hand, several limitations should be considered when interpreting the results presented in this study. First, we examined only acute effects; whether these changes persist or accumulate with repeated stimulation remains unknown. Second, the absence of physiological measures (e.g., heart rate variability, pupillometry, or salivary biomarkers) limits our ability to confirm proposed mechanistic pathways. Finally, given the no statistical significant results from ultrasound stimulation when comparing active and sham conditions, but the statistical significant results between pre- and post-stimulation measures for active, together with the similar trajectory of the results on its sham condition (as shown in Fig. 3), which are opposed to sham E-taVNS, it can be concluded that the auditory masking of U-taVNS might be having an unexpected impact on working memory performance. This suggests that a better masking mechanism or control group might be necessary to better validate if the similar but not significant performance of U-taVNS compared to E-taVNS could be due to such sham condition rather than a different in efficacy between both modalities.

Furthermore, given the lower aversive effects of U-taVNS compared to E-taVNS, further research with a bigger population group, other sham comparison conditions, and other protocol parameters is necessary to further conclude if ultrasound stimulation through taVNS is as effective as electrical stimulation for working memory, or if the intensity of the stimulation needs to be increased and the sham condition better controlled.

Despite these limitations, the findings carry meaningful clinical implications. Demonstrating that both taVNS modalities can improve working memory performance in healthy individuals provides proof-of-concept for potential therapeutic applications in neurodegenerative disorders. The superior tolerability of U-taVNS, even with a reduced effect compared to its own sham, positions it as a promising candidate for home-based and long-term therapeutic protocols.

## Conclusion

In conclusion, this study provides evidence that electrical-based taVNS can acutely modulate working memory performance in healthy adults, while ultrasound-based taVNS shows a similar trend to electrical stimulation. Although these findings advance our understanding of non-invasive Vagus Nerve stimulation as a tool for modulating cognitive processing, further research is necessary to better understand the effects of ultrasound-based stimulation of the Vagus Nerve for cognitive processes. As the field moves toward clinical implementation, our results support continued investigation of taVNS approaches for treating conditions characterized by neurodegenerative outcomes.

## Abbreviations

E-taVNS: electrical transcutaneous auricular Vagus nerve stimulation
U-taVNS: ultrasound transcutaneous auricular Vagus nerve stimulation
WM: working memory.

## Acknowledgments

The authors acknowledge all participants.

