## Supplementary Material for "Transcutaneous auricular Vagus nerve stimulation for working memory enhancement: A comparative study of electrical and ultrasound stimulation"

### Supplementary Materials

#### Pre- and Post-stimulation performance Metrics

Results showed that there was a significant main effect of Phase (Pre vs Post, Correct rejection: $t$ (30) = 2.98, $p$ = 0.006, Cohen's $d$ = 0.54; Error False alarm: $t$ (30) = −2.98, $p$ = 0.006, Cohen's $d$ = −0.54) on active E-taVNS and U-taVNS (Overall accuracy: $t$ (28) = 2.23, $p$ = 0.034, Cohen's $d$ = 0.42; $d'$: $t$ (28) = 2.72, $p$ = 0.012, Cohen's $d$ = 0.51), and sham E-taVNS (Correct rejection: $t$ (30) = −3.52, $p$ = 0.001, Cohen's $d$ = −0.64; Error False alarm: $t$ (30) = 3.52, $p$ = 0.001, Cohen's $d$ = 0.64; $d'$: $t$ (30) = −2.11, $p$ = 0.044, Cohen's $d$ = −0.38) reflecting changes across pre- and post-stimulation assessments, as shown in Table S1.

When analyzing baseline differences between modalities, we found statistically significant difference in active stimulation baseline for average hit reaction time ($t$ (58) = −2.01, $p$ = 0.049, Cohen's $d$ = 0.53), but not significant difference in sham stimulation baseline between E-taVNS and U-taVNS. Thus, we used the Post/Pre ratio metrics for following analyses.

**Table S1.** Pre- and post-stimulation performance metrics for E-taVNS and U-taVNS. Values represent mean ± standard deviation (SD). Significance for the Pre vs Post paired $t$-test is indicated by bold $p$-values ($p$ < 0.05). Hit RT represents the average Hit reaction time, False Alarm represents the error false alarm, and Correct Reject represents correct rejection rate.

| **Modality** | **Metric** | **Condition** | **Mean ± SD** | | ***t* (df=58)** | ***p*** | **Cohen's *d*** |
|---|---|---|---|---|---|---|---|
| | | | Pre | Post | | | |
| **E-taVNS** | Overall Accuracy | Active | 0.93 ± 0.07 | 0.95 ± 0.06 | 1.90 | 0.068 | 0.35 |
| | | Sham | 0.95 ± 0.05 | 0.94 ± 0.04 | -0.83 | 0.414 | -0.15 |
| | Hit Accuracy | Active | 0.88 ± 0.14 | 0.84 ± 0.18 | -1.42 | 0.166 | -0.26 |

| | | | | | | | |
|---|---|---|---|---|---|---|---|
| | | Sham | 0.86 ± 0.14 | 0.88 ± 0.13 | 0.86 | 0.399 | 0.16 |
| | Correct Rejection | Active | 0.95 ± 0.09 | 0.99 ± 0.02 | 2.98 | **0.006** | 0.54 |
| | | Sham | 0.98 ± 0.03 | 0.96 ± 0.02 | -3.52 | **0.001** | -0.64 |
| | Error Miss | Active | 0.12 ± 0.14 | 0.16 ± 0.18 | 1.42 | 0.166 | 0.26 |
| | | Sham | 0.14 ± 0.14 | 0.12 ± 0.13 | -0.86 | 0.399 | -0.16 |
| | False Alarm | Active | 0.05 ± 0.09 | 0.01 ± 0.01 | -2.98 | **0.006** | -0.54 |
| | | Sham | 0.02 ± 0.03 | 0.04 ± 0.02 | 3.52 | **0.001** | 0.64 |
| | Hit RT | Active | 0.78 ± 0.19 | 0.78 ± 0.20 | 0.20 | 0.841 | 0.04 |
| | | Sham | 0.80 ± 0.23 | 0.78 ± 0.19 | -0.87 | 0.394 | -0.16 |
| | *d'* | Active | 2.87 ± 0.65 | 3.09 ± 0.71 | 1.86 | 0.073 | 0.34 |
| | | Sham | 3.13 ± 0.71 | 2.89 ± 0.54 | -2.11 | **0.044** | -0.38 |
| **U-taVNS** | Overall Accuracy | Active | 0.92 ± 0.06 | 0.94 ± 0.06 | 2.23 | **0.034** | 0.42 |
| | | Sham | 0.94 ± 0.05 | 0.95 ± 0.05 | 1.53 | 0.136 | 0.29 |
| | Hit Accuracy | Active | 0.84 ± 0.13 | 0.88 ± 0.15 | 1.46 | 0.157 | 0.28 |
| | | Sham | 0.86 ± 0.10 | 0.90 ± 0.12 | 1.60 | 0.121 | 0.30 |
| | Correct Rejection | Active | 0.96 ± 0.07 | 0.97 ± 0.06 | 1.92 | 0.066 | 0.36 |
| | | Sham | 0.096 ± 0.06 | 0.97 ± 0.05 | 0.69 | 0.496 | 0.13 |
| | Error Miss | Active | 0.16 ± 0.13 | 0.12 ± 0.16 | -1.46 | 0.157 | -0.28 |
| | | Sham | 0.14 ± 0.10 | 0.10 ± 0.12 | -1.60 | 0.121 | -0.30 |
| | Error False Alarm | Active | 0.04 ± 0.07 | 0.03 ± 0.06 | -1.92 | 0.067 | -.036 |
| | | Sham | 0.04 ± 0.06 | 0.03 ± 0.06 | -0.69 | 0.496 | -0.13 |
| | Hit RT | Active | 0.86 ± 0.17 | 0.86 ± 0.17 | -0.29 | 0.776 | -0.05 |
| | | Sham | 0.884 ± 0.17 | 0.84 ± 0.20 | -1.48 | 0.151 | -0.28 |
| | *d'* | Active | 2.76 ± 0.65 | 3.13 ± 0.75 | 2.72 | **0.012** | 0.51 |
| | | Sham | 2.91 ± 0.64 | 3.13 ± 0.49 | 1.58 | 0.125 | 0.30 |

### Baseline Questionnaires as Outcome Predictors

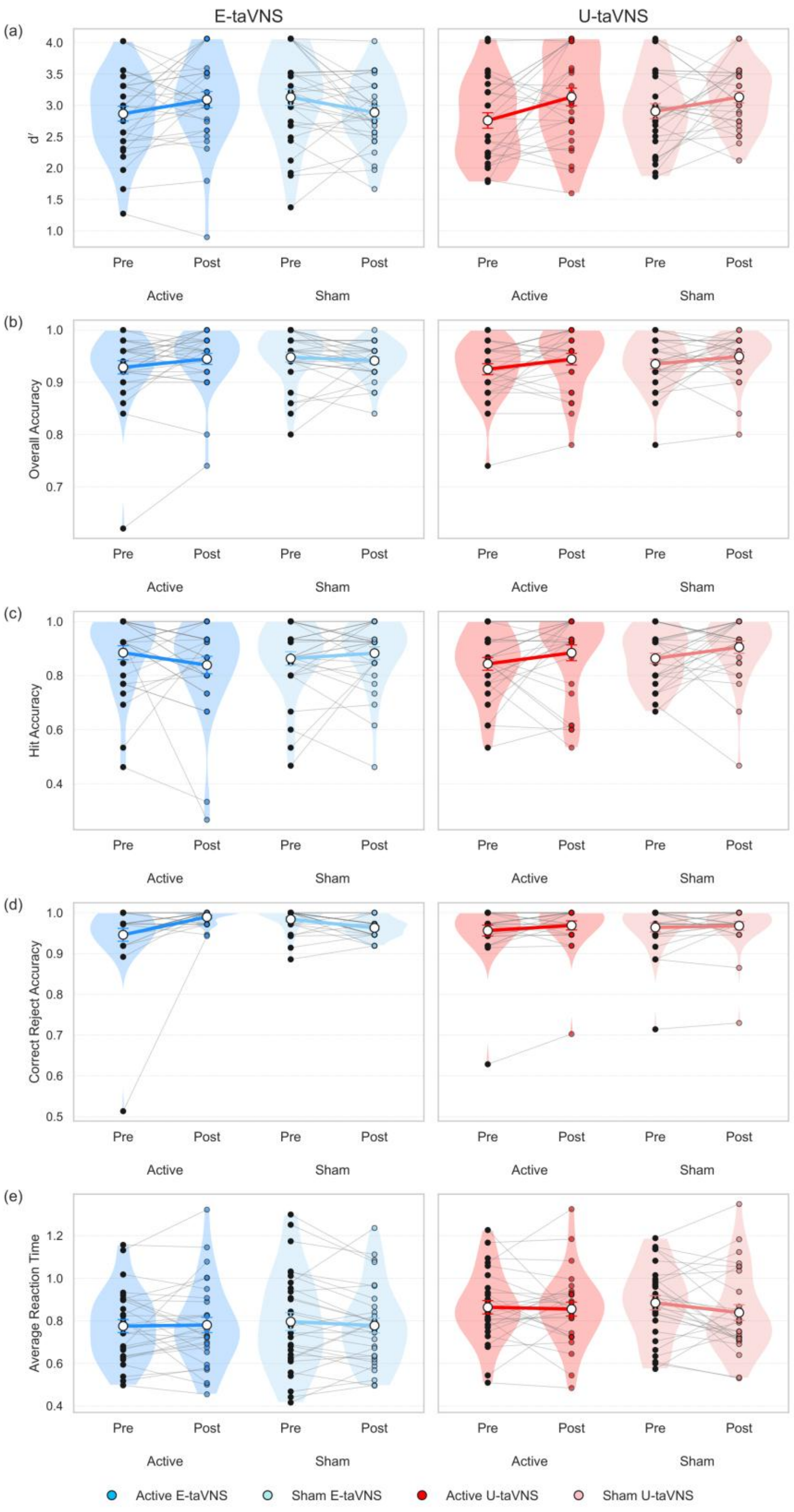


**Figure S1.** Individual and group-level changes in working memory performance from pre- to post-stimulation. Pre- to post-stimulation values in working memory metrics are shown for E-taVNS and U-taVNS under active and sham conditions. Each panel (a-e) corresponds to a different metric (*d'*, Overall Accuracy, Hit Accuracy, Correct Rejection rate, and average Hit Reaction Time), with E-taVNS results represented at the left side and U-taVNS at the right side of each panel. Thin grey lines represent individual participant trajectories from pre- to post-stimulation, with colored points indicating pre- and post-stimulation values for active and sham conditions. Thick colored lines depict group means, with error bars indicating SD. Colors denote stimulation condition within each modality (Active vs Sham), allowing visual comparison of stimulation-specific effects within and across modalities.

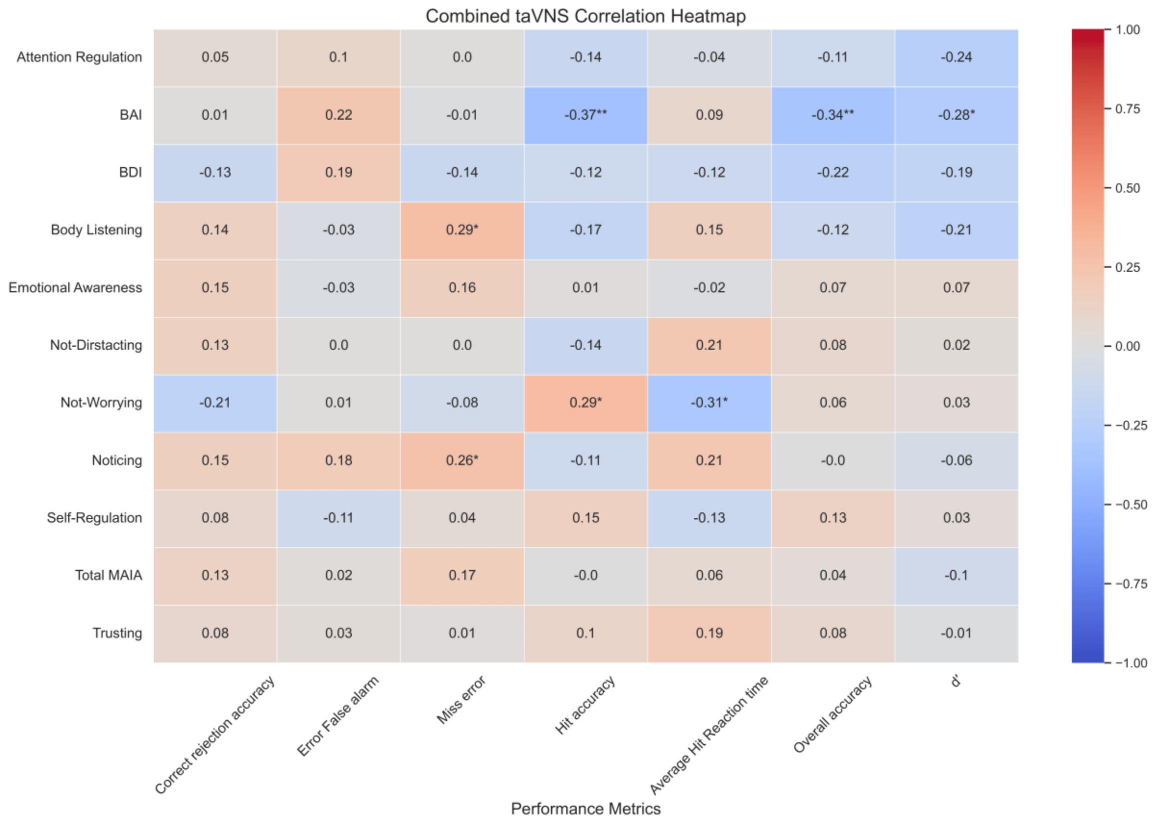


**Figure S2.** Relationship between Questionnaire scores and outcome of combined taVNS in working memory performance as measured by ratio changes (Post/Pre Active/Post/Pre Sham). Figure depicts the Pearson correlation heatmap between baseline questionnaire scores and the combined taVNS. Statistical significance is represented by ‘*’ for $p < 0.05$, and ‘**’ for $p < 0.01$.

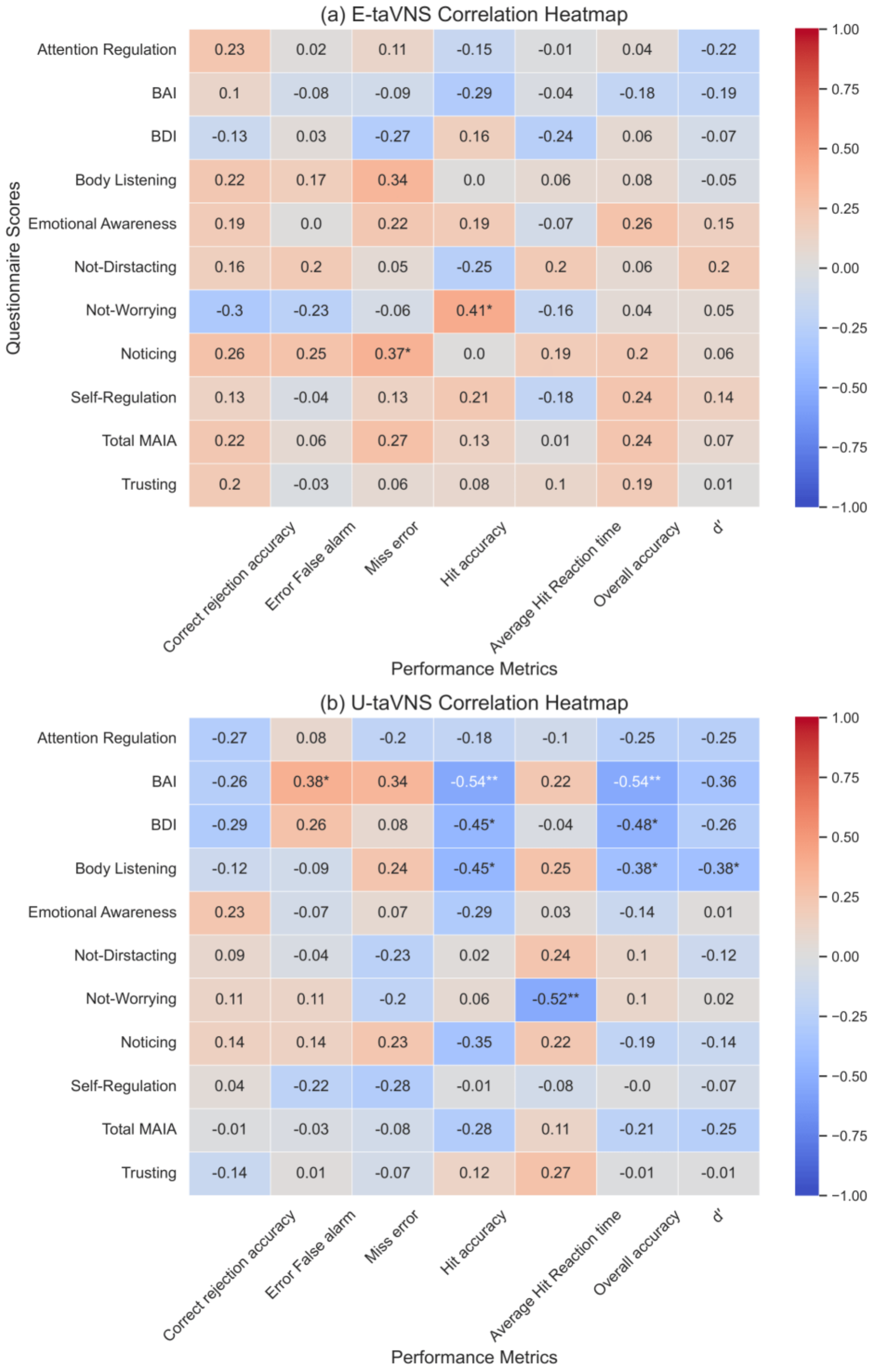


**Figure S3.** Relationship between questionnaire scores and outcome of taVNS in working memory performance as measured by ratio changes (Post/Pre

Active/Post/Pre sham) per modality. The figure depicts the Pearson correlation heatmap between baseline questionnaire scores and (a) E-taVNS, and (b) U-taVNS. Statistical significance is represented by '*' for $p < 0.05$, and '**' for $p < 0.01$. U-taVNS is more influenced by baseline differences than E-taVNS.

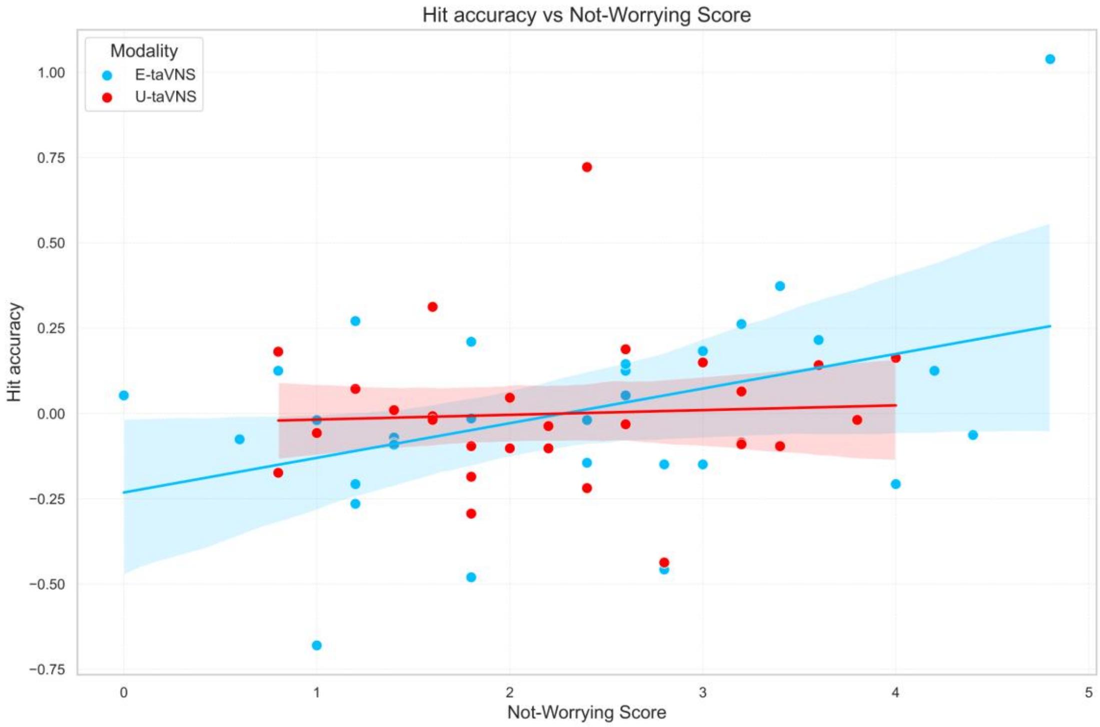


**Figure S4.** Relationship between Not-worrying scores and outcome of taVNS in working memory performance for Hit Accuracy. Scatterplot illustrates the association between Not-worrying scores and working memory performance after taVNS, as measured by ratio changes (active/sham) in Hit Accuracy. Each point represents individual participants. Blue dots indicate participants that received E-taVNS, while red dots represent participant that received U-taVNS. A higher Not-worrying score was associated with a better performance and a higher effect of the stimulation. E-taVNS: B = 0.102, SE = 0.043, t = 2.35, $p = 0.026$; U-taVNS: B = 0.014, SE = 0.046, t = 0.30, $p = 0.766$.

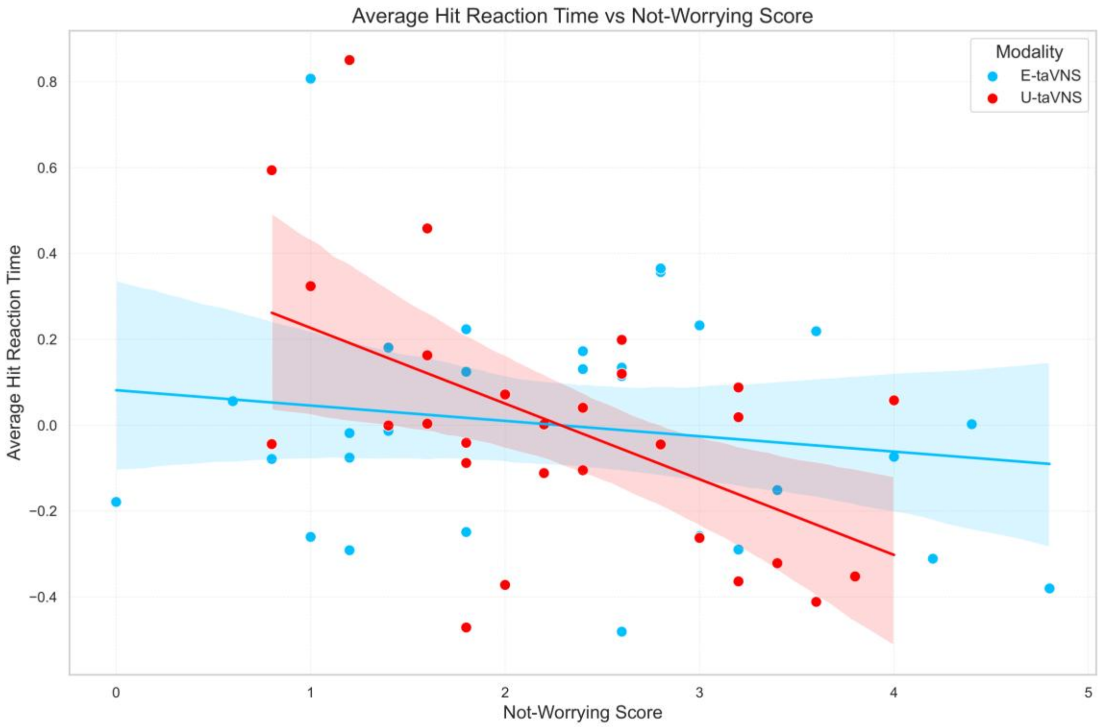


**Figure S5.** Relationship between Not-worrying scores and outcome of taVNS in working memory performance for Average Hit Reaction Time. Scatterplot illustrates the association between Not-worrying scores and working memory performance after taVNS, as measured by ratio changes (active/sham) in Average hit reaction time. Each point represents individual participants. Blue dots indicate participants that received E-taVNS, while red dots represent participant that received U-taVNS. A higher Not-worrying score was associated with a better performance through a smaller reaction time and a higher effect of the stimulation. E-taVNS: B = -0.036, SE = 0.041, t = -0.87, $p$ = 0.392; U-taVNS: B = -0.176, SE = 0.056, t = -3.15, $p$ = 0.004.
